# Alternative promoters in CpG depleted regions pervasively account for epigenetic misregulation of cancer transcriptomes

**DOI:** 10.1101/2022.08.02.502575

**Authors:** Chirag Nepal, Jesper B. Andersen

## Abstract

Eukaryotic genes are regulated by multiple alternative promoters with distinct expression patterns. In cancer, alternative promoters are pervasively utilized, but our understanding of the mechanism of activation and how their regulatory architecture differs from reference promoters remains elusive. We analyzed 100 CAGE-seq libraries from HCC patients and annotated 4083 alternative promoters in 2926 multi-promoter genes that are known genes involved in hepatocarcinogenesis. Many alternative promoters are undetected in the normal liver. We find that multi-promoter genes are enriched among genes downregulated in the tumor, highlighting alternative promoters’ impact in global transcription changes in cancer. Alternative promoters are depleted for CpG islands, have narrow nucleosome depleted regions, and are enriched for sharp promoters as well as tissue-specific transcription factors. Alternative promoters have high DNA methylation levels around transcription start sites. Tumor cells globally lose DNA methylation, but there exists a hierarchical retention of intragenic DNA methylation, which is dictated by the genomic CG content. As such, intragenic CG-poor regions lose methylation, while CG-rich regions retain it, a phenomenon caused by differential binding of H3K36me3, *DNMT3B, TET1* and *SETD2.* Thus, the selective loss of DNA methylation in CG-poor regions opens the chromatin and makes these regions accessible for transcription. Upon transcription factors availability, alternative transcription can pervasively occur in cancer. These results provide a framework for understanding the importance of alternative promoters in controlling the tumor transcriptomes, highlighting their architecture and role in regulatory mechanism(s).

## Introduction

Hepatocellular carcinoma (HCC) accounts for about 90% of primary liver cancers and is the fourth most common cause of cancer-related death^1^. Genome-wide profiling of HCC patients has helped to build a molecular map of mutations, dysregulated genes, and DNA methylomes^2–5^. Additional efforts to map histone modifications^6^, chromatin accessibility^7^, transposon activation^8^ and RNA m6A methylation^9^ are ongoing and important to understand mechanisms of gene regulation in cancer. Promoters are gateways to start transcription and regulate gene expression in a temporal and spatial manner. Yet, there has been little effort to understand promoter regulation in cancer. In this context, how the regulatory architecture of promoter usage impacts gene regulation in HCC is unclear.

Transcription is facilitated by a reference promoter, which is the region proximal to the transcription start site (TSS), integrating *cis*-regulatory elements to ensure precise gene regulation^10^. Cap Analysis of Gene Expression sequencing (CAGE-seq) determines the 5’-ends of TSSs at single nucleotide resolution^11,12^ and quantifies gene expression similar to RNA-seq^13^. CAGE-seq also detects differently regulated transcription initiation events within the same core promoter^14,15^, and thus, it can accurately detect alternative promoters. In addition to reference promoters, many genes utilize alternative promoters for specific processes such as cell fate transitions in yeast^16^ and mammalian cells^17^, in vertebrate embryogenesis^12,18^, and are believed to have important roles in cancers^19^. Widespread activation of alternative promoters is known in different contexts; however, it is unclear whether their (epi-)genetic states are different compared to the reference promoter, and if they are under a different regulatory architecture. Alternative promoters have dynamic intragenic DNA methylation across human tissues^20^ and loss of DNA methyltransferase 3B (DNMT3B) in mouse embryonic stem (ES) cells has been shown to result in spurious initiation of alternative promoters^21^. Therefore, precise regulation of alternative promoters is important to ensure correct gene expression.

Alternative promoters are pervasively activated in cancer^19^. Our understanding of the mechanism(s) of activation and their impact on gene expression is unknown. We analyzed 100 CAGE-seq libraries from HCC patients and comprehensively annotated 4083 alternative promoters, representing 2926 multi-promoter (MP) genes, supported by histone modifications, ATAC-seq, RNA Pol2 Chip and DNA methylation across the patient cohort and HepG2 cells. Transcription of alternative promoters is dominant outside CpG islands, enriched for genes important in HCC, and their activation frequently results in downregulation of the reference promoter. We showed that CG-poor regions preferentially loose DNA methylation in tumors, followed by chromatin opening and Pol2 binding, resulting in alternative transcription from CG-poor regions. Collectively, our study elucidates the mechanism of activation and preferential underlying DNA sequence for alternative promoter activation in cancer.

## Results

### Pervasive transcription of alternative promoters in HCC

To determine the extent of alternative promoter usage in HCC, we analyzed CAGE-seq data from 50 tumors and their matched tumor-adjacent tissues^8^ (Supplementary Table 1). We identified the 5’-end of CAGE transcription start sites (CTSSs) and quantified expression levels in tags per million (TPM). Proximal CTSSs within 20 nucleotides on the same strand were clustered to define transcript clusters (TCs) (Fig. 1A and Supplementary Fig. 1A-B). We retained TCs expressed above 1 TPM in at least 15 samples and a minimum 3 TPM in at least one sample, which resulted in 42804 high-confidence consensus TCs expressed across the cohort (Fig. 1A and Supplementary Table 2). A majority of TCs (90%) were supported by FANTOM5 CAGE peaks^17^ and/or open chromatin regions from ENCODE^22^ and TCGA^7^ (Fig. 1B). Based on GENCODE transcript models, we identified promoters for 15419 expressed genes and alternative TSSs for 3052 annotated alternative transcripts. A significant fraction (10492; 24.5%) of CAGE TCs were in intragenic regions (Fig. 1C), indicating these are putative alternative promoters. We filtered intragenic CAGE TCs that represent drosha processing of pre-miRNAs^23^, snoRNAs 5’-ends capping^17^, exons post-transcriptional processing^11,12^, enhancer RNAs^24^ and promoters lacking a transcription initiator^15^ (Fig. 1C). For the remaining TCs, we clustered all within 300 bases, resulting in 1031 novel alternative TSSs (Supplementary Table 3). A majority of these novel TSSs were supported by RNA-seq and expressed sequence tag transcripts (Fig. 1D). In total, we identified 4083 alternative TSSs in HCC (represented by 3052 annotated TSSs and 1031 novel TSSs) (Fig. 1C).

**Figure 1.**
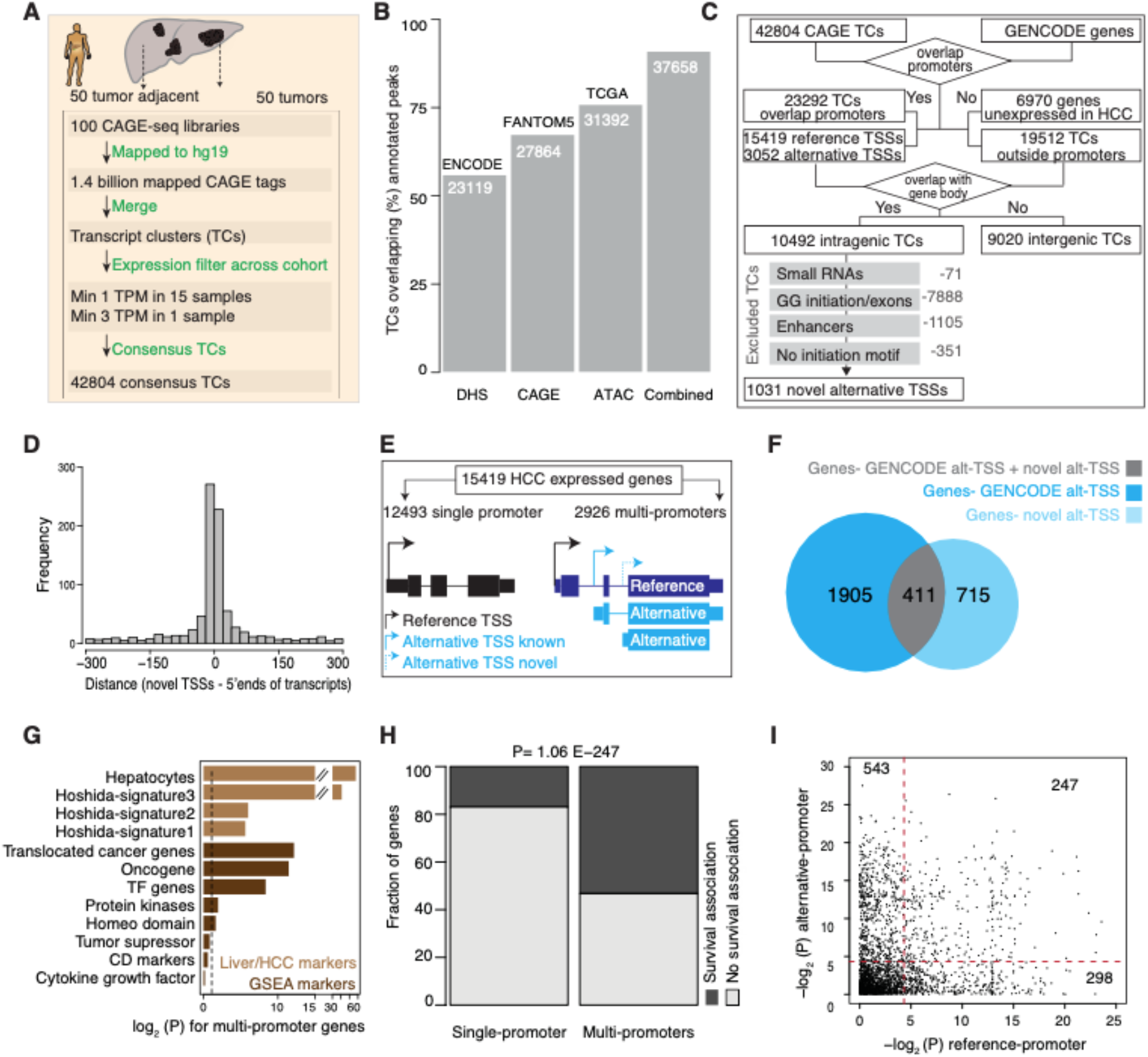
Annotation of alternative promoters in hepatocellular carcinoma (HCC) patients. (**A**) A schematic workflow to describe the mapping of CAGE-seq reads to define consensus transcript clusters (TCs) across the cohort. (**B**) Barplot shows overlap of HCC TCs with the annotated FANTOM5 CAGE peaks and open chromatin peaks from ENCODE and TCGA. (**C**) A schematic workflow to annotate intragenic CAGE TCs as high-confidence alternative promoters. The workflow includes multiple filtering steps to exclude TCs that lack promoter features. (**D**) Distance between 5’ ends of novel TSSs and 5’ ends of RNA-seq and EST transcripts. (**E**) Classification of expressed genes into single promoter (SP) and multi-promoter (MP) genes based on the number of promoters. The promoter with highest expression level (represented by arrow height) is assigned as the reference promoter. (**F**) Venn diagram shows the intersection of novel alternative promoters with known MP genes. (**G**) Enrichment of signature genes in MP genes compared to SP genes. (**H**) Distribution of survival associated genes with SP and MP genes. The MP genes were significantly associated (P= 1.06E-247; Fisher’s exact test) with survival outcome. (**I**) The scatter plot shows the p-value of the survival association for reference and alternative promoters.

Based on the number of promoters, we classified 15419 liver expressed genes into either single-promoter (SP) (12493;80.3%) or multi-promoter (MP) (2926;19.7%) genes (Fig. 1E, Supplementary Table 3 and Supplementary Table 4). Promoters with the highest mean expression level across the cohort were assigned as the reference (major) promoter, whereas the others were assigned as alternative (minor) promoters (Fig. 1E, see methods). Most novel alternative TSSs occurred in genes without active alternative promoters (Fig. 1F), while others occurred in existing MP genes, resulting in some genes with multiple alternative promoters (Supplementary Fig. 1C). Overall, 65% of alternative promoters are located downstream of the reference promoter (Fig. 1G) and 35% are upstream, which highlights that purely assigning the most upstream TSS as the reference promoter not always is optimal. Alternative promoters utilized the same N terminus in 986 genes and different N terminus of the proteins in 1819 genes (Supplementary Table 3), as exemplified for *CDKN2A* (Supplementary Table 3) and *ERBB2* (Supplementary Fig. 1E). MP genes were associated with diverse functions enriched in metabolic processes, signalling pathways, apoptosis, regulation of cell migration and programmed cell death (Fig. S1F and Supplementary Table 5), which is in contrast with SP genes that are overrepresented by housekeeping functions (translation, DNA repair, gene expression, mRNA splicing). Notably, hepatocytic markers^25^, HCC signature genes^26^ and cancer-associated gene families (oncogenes, transcription factors and protein kinases) from GSEA^27^ were over-represented in MP genes (Fig. 1G). Next, we reanalyzed TCGA HCC clinical data and observed a significant association (p=3.2E-78; Fisher’s exact test) of survival-gene expression among MP genes compared to SP genes (Fig. 1H and Supplementary Table 3). Compared to reference promoters, the expression of alternative promoters was superior in predicting overall survival of HCC patients (Fig. 1J). This includes both genes known/unknown to HCC, and genes not previously described to rely on promoter switching, such as *SULF2* (Supplementary Fig. 1G). Alternative promoters with survival-association had shorter overall survival time (Supplementary Fig. 1H). Moreover, MP genes were enriched for sorafenib resistance marker-genes^28^ (Supplementary Fig. 1I). Collectively, these data support that alternative promoters and their usage are important in the HCC pathogenesis and for patient’s outcome.

### Impact of alternative promoters on gene expression

We sought to understand how alternative promoters control gene expression. We first determined the fraction of HCC reference and alternative promoters that are expressed across 8 independent normal livers^8,17^. Whereas, some HCC alternative promoters were undetectable in the normal liver, they are already expressed in the matched tumor-adjacent liver (cirrhotic liver parenchyma) (Fig. 2A, Supplementary Fig. 2A and Supplementary Table 4). At the expression threshold of minimum 3 TPM in at least 1 normal liver sample approximately 50% of the alternative promoters were unexpressed in the normal liver compared to 15% of reference promoters (Fig. 2B). We lowered the expression threshold and observed a higher fraction of alternative promoters were undetected in the normal livers. MP genes with alternative promoters expressed in normal livers were enriched for metabolic-related pathways, reflecting a tissue-intrinsic biology. On the contrary, tumor-specific alternative promoter genes were enriched in oncogenic pathways such as, WNT/beta-catenin signalling, E2F and Myc targets (Fig. 2C and Supplementary Table 6).

**Figure 2.**
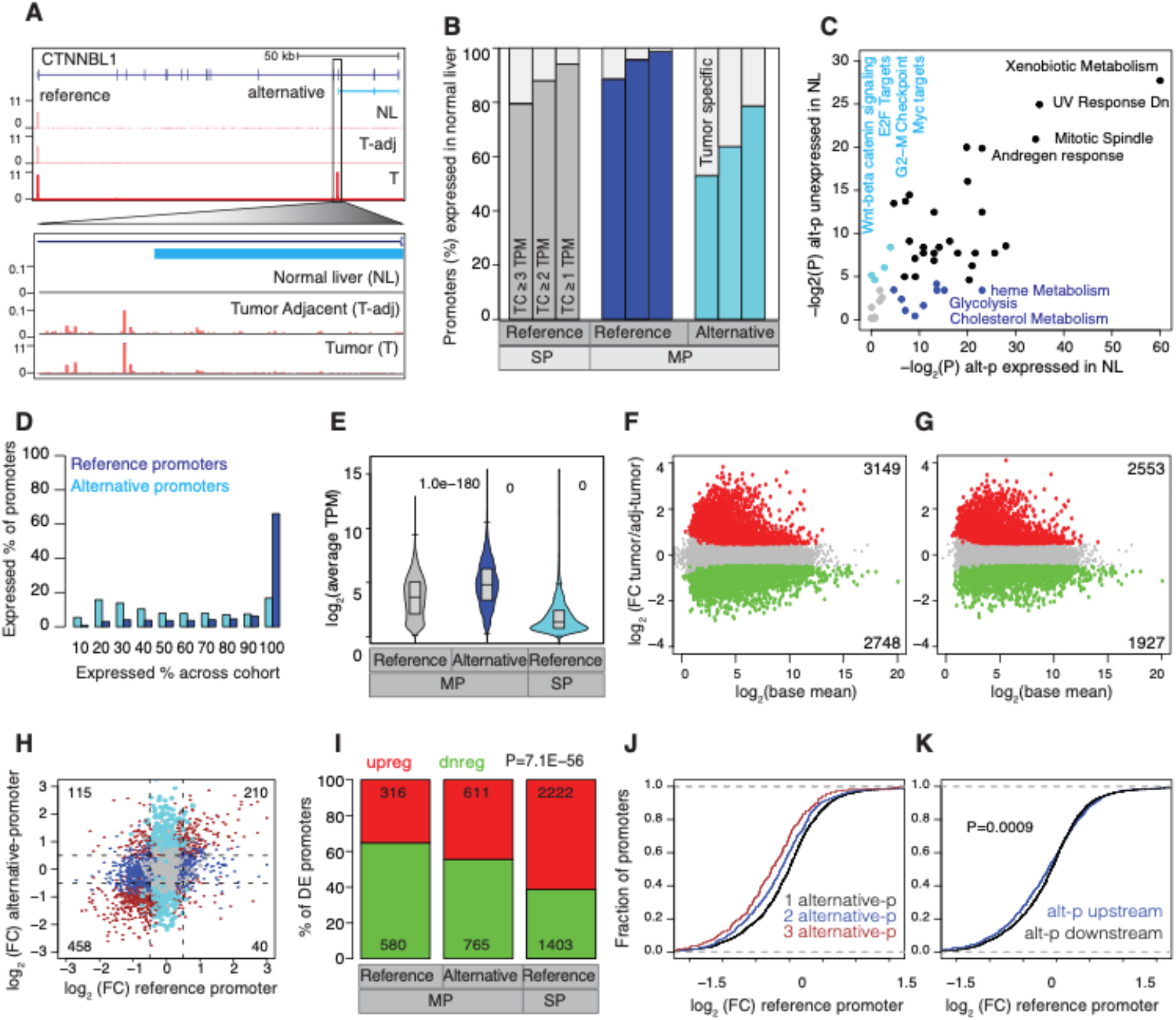
The impact of alternative promoters in gene expression regulation. (**A**) A UCSC browser screenshot of the *CTNNBL1* gene with CAGE tags across the normal liver, tumor-adjacent tissues, and HCCs. The alternative promoter is zoomed in to show the CTSS usage. (**B**) Barplot shows the percentage of HCC promoters that are transcribed across 8 independent normal livers at different threshold of CAGE TCs. HCC Promoters undetected at low levels show tumor-specific activation of alternative promoters. Expressed genes are classified into single promoter (SP) and multi-promoter (MP) genes, where MP genes are further classified into reference and alternative promoters. (**C**) Enriched cancer hallmark terms associated with alternative promoters expressed/unexpressed in normal livers. (**D**) Distribution of reference and alternative promoters expressed across the HCC cohort. The x-axis indicates the percentage, at which a promoter is expressed across the HCC cohort. (**E**) The average expression level of reference and alternative promoters of multi-promoter genes and single promoter genes. (**F-G**) Volcano plots show differentially expressed promoters (F) and genes (G) between tumors and tumor-adjacent tissues. The cut-off p-value of 0.05 was FDR-corrected. (**H**) Expression fold change for reference and alternative promoter pairs. (**I**) Barplot shows the fraction of differentially expressed promoters (from panel F) that are classified as either upregulated or downregulated in tumors compared to tumor-adjacent tissues. (**J-K**) The cumulative plots show the distribution of fold change of reference promoters compared to the number of alternative promoters (F) or based on the position (upstream/downstream) of the alternative promoter relative to the reference promoter.

We computed the extent of reference and alternative promoters expressed across the patient cohort and showed that the reference promoters (irrespective of SP or MP genes) were constitutively expressed across the cohort, while alternative promoters were either constitutively expressed or expressed in a subset of patients (Fig. 2D). Notably, alternative promoters unexpressed in normal livers generally were expressed in a subset of HCCs (Supplementary Fig. 2B). Although expression levels of alternative promoters on average were lower (Fig. 2E) at the population level some alternative promoters were higher than its reference promoters at the individual patient level (Fig. S2A). In tumor-adjacent tissues, we identified 1489 (36.4%) alternative promoters with higher expression than the reference promoters in minimum one patient which further increased to 1716 (42%) alternative promoters when analyzing tumors alone (Supplementary Fig. 2C). In fact, the variance in the expression levels were highest for alternative promoters across the cohort (Supplementary Fig. 2D).

In total, we identified 3200 promoters and 2400 genes differentially expressed^29^ between tumor and tumor-adjacent tissues (Fig. 2F-G and Supplementary Table 7). Almost all (99%) of the differentially expressed genes were detected at the promoter level (Supplementary Fig. 2E). We compared the expression fold-change of reference and alternative promoters among MP genes and observed a general trend of downregulation of reference promoters and upregulation of alternative promoters (Fig. 2H). In total, we identified 155 promoter pairs as significantly different (log2 fold-change of 0.5) with expression in opposite direction, among which 115 (74%) of the promoter pairs the reference promoters were downregulated and alternative promoters were upregulated (Fig. 2H). It has been shown that alternative upstream promoters^30^ or downstream intragenic promoters^31^ attenuate host gene expression, thus we asked whether genes downregulated in HCC are enriched among MP genes. Among the differentially expressed promoters, MP genes were mostly downregulated, while SP genes were mostly upregulated (Fig. 2I), which held true across different functional classification of genes except for KEGG signaling pathways (Supplementary Fig. 2F). Indeed, upstream promoters are known to interfere with downstream reference promoters^30^ and downstream intragenic promoters attenuate host gene expression^31^. Consistent with this observation, we showed that the reference promoters were downregulated when alternative promoters are located upstream (Fig. 2J). Downregulation of reference promoters was more evident when a gene has more than one alternative promoter (Fig. 2K), which may interfere with the elongating polymerase leading to transcript downregulation. Collectively, this highlights that alternative promoters are pervasively transcribed in tumors affecting many key cancer-related pathways and often leading to downregulation of reference promoters.

### Alternative promoters have distinct underlying DNA sequence and promoter architecture

Two thirds of human genes have CpG islands (CGIs) within their promoters^32^ as depicted for examples *GNAS* (Fig. 3A), *ERBB2* (Fig. 3B) and *COMT* (Supplementary Fig. 3A). In addition, there are thousands of intragenic CGIs (not associated with known promoters) that act as alternative promoters during development^33^, prompting us to ask whether intragenic CGIs function as alternative promoter in HCC. Overall, 80% of reference promoters overlap with CGIs, whereas this is 45% (1899 out of 4083) for alternative promoters (Fig. 3C), reflecting the depletion of CG dinucleotides in alternative promoters (Supplementary Fig. 3B). Furthermore, in a significant fraction (981 out of 1899) the CGIs were shared between the alternative and reference promoters (Fig. 3A and Fig. 3C). The shared CGIs are the longest, while those exclusive to alternative promoters have the shortest length (Supplementary Fig. 3C). Also, compared to known alternative promoters, CGIs were further depleted among novel alternative promoters (Fig. 3D), reflecting a preferential activation of novel promoters from CG-poor regions.

**Figure 3.**
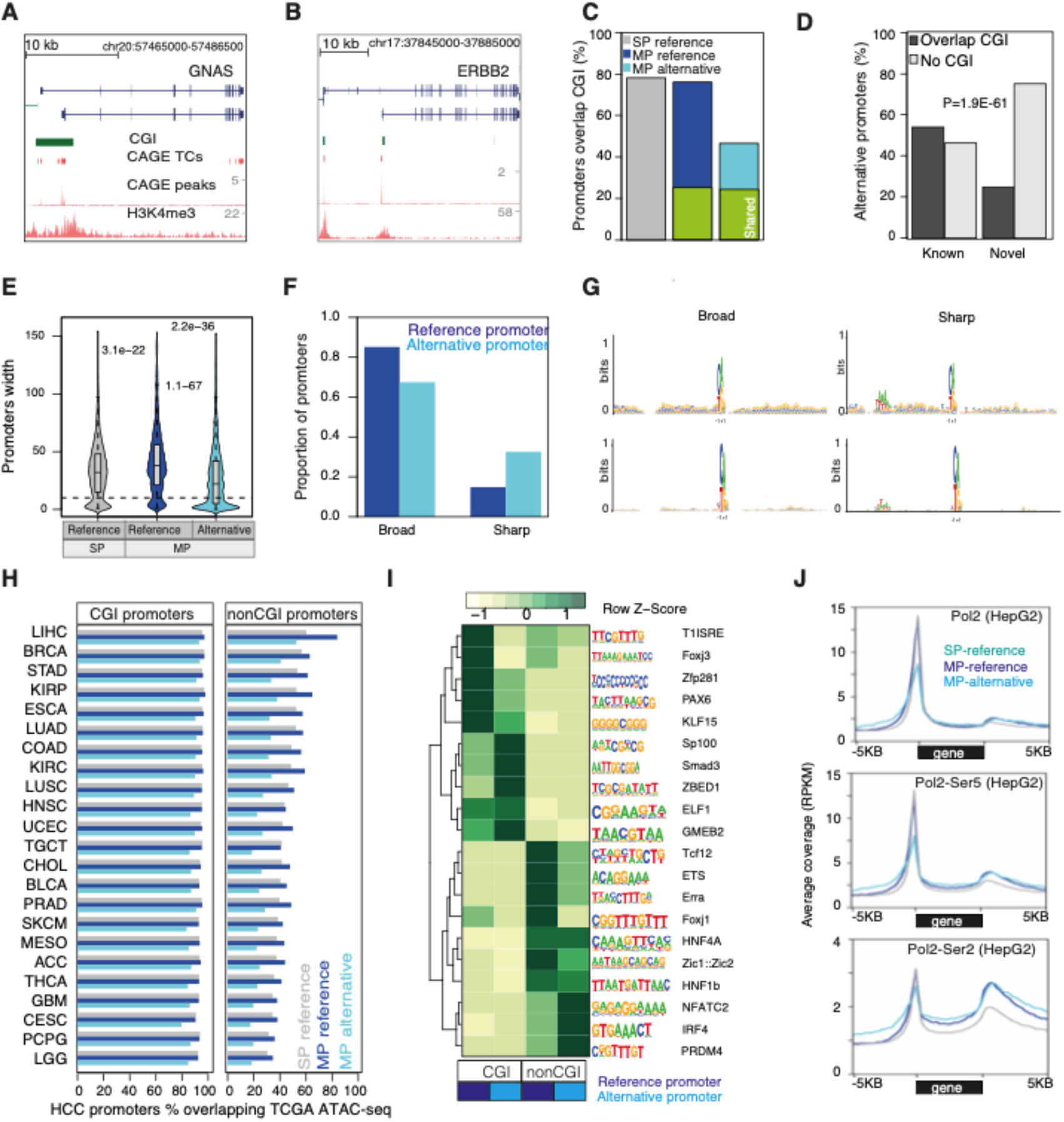
Promoter architecture, sequence composition and motifs of reference and alternative promoters. (**A-B**) A UCSC browser screenshot of *GNAS* (**A**) and *ERBB2* (**B**) genes along with CpG islands (CGI), CAGE-seq and H3K4me3 tracks. The reference and alternative promoters of *GNAS* have a shared CGI. The reference and alternative promoters of *ERBB2* have distinct nonoverlapping CGIs. (**C**) Barplot shows the fraction of expressed reference and alternative promoters that overlapped with CGIs. CGIs shared by reference and alternative promoters are highlighted in green. SP and MP denote single-promoter and multi-promoter genes, respectively. (**D**) Barplot shows the fraction of annotated and novel alternative promoters overlapping with CGIs. (**E**) Boxplot shows the distribution of promoter width. (**F**) Barplot shows alternative promoters have a higher proportion of sharp promoter shape relative to reference promoters. (**G**) Different sequence motifs around TSSs of sharp and broad promoters both for reference and alternative promoters. (**H**) Barplot shows the fraction of HCC promoters that overlapped with TCGA pan-cancer ATAC-seq peaks. Promoters were classified into two groups based on their overlap with CGIs. (**I**) Heatmap shows transcription factor motifs enriched across reference and alternative promoters as well as their overlap with CGIs. (**J**) The average coverage of RNA polymerase II (Pol2), Pol2-Ser5 (initiation of Pol2) and Pol2-Ser2 (elongation by Pol2) in HepG2 cells across the gene body of SP and MP genes.

We hypothesized that to adapt to the evolving cancer transcriptome, alternative promoters might prefer a distinct regulation than the reference promoter, which might be mediated via promoter architecture, chromatin accessibility, transcription factor-association and RNA Pol2 dynamics. Reference promoters have significantly broader promoter width (broad promoters) (Fig. 3E-F), while alternative promoters are narrow and associated with sharp promoters (Fig. 3F). Sharp promoters have one dominant transcription initiation site, while broad promoters have multiple initiation sites with a transcript cluster (Supplementary Fig. 3D-E). Sequence alignment around TSSs revealed an initiator motif across all promoters, while the TATA box was positionally constrained upstream of sharp promoters (Fig. 3G), as previously observed^11,12^. We analyzed chromatin accessibility of HCC promoters across TCGA ATAC-seq peaks^7^ and expectedly showed the highest overlap with HCC (Fig. 3H). Majority of reference and alternative CGI promoters were uniformly accessible across cancers while chromatin accessibility of nonCGI promoters varied across tumor types (Fig. 3H). Thus, CG-rich reference promoters have ubiquitous expression while CG-poor alternative promoters have tissue-specific expression. ENCODE transcription factor ChIP-seq peaks were significantly higher among reference promoters (Supplementary Fig. 3F). *De novo* analysis of transcription factors revealed that reference and alternative nonCGI promoters were enriched for liver-specific transcription factors (Supplementary Table 8) such as hepatocyte nuclear factor 1A (HNF1A) and HNF4B (Fig. 3I), which is consistent with previously observed tissue-specific transcription factor enrichment among nonCGI promoters^34^. Notably, the transcription factor NFATC2 was enriched only on nonCGI alternative promoters. Recently, hepatic NFAT signaling was reported to regulate inflammatory cytokine expression in cholestasis^35^. Lastly, RNA polymerase II (Pol2) was enriched at similar levels in both SP and MP genes, but both initiation (Pol2-Ser5) and elongation (Pol2-Ser2) by Pol2 were higher among MP genes (Fig. 2J). These data were validated by GRO-seq (Supplementary Fig. 3G). Thus, alternative promoters have distinct regulatory architecture characterized by low CG content, sharp promoter shape and enriched for tissuespecific transcription factors.

### Alternative promoters have distinct histone modifications and nucleosome positioning

Having established that most alternative promoters are CG-poor, we sought to understand whether this intrinsic sequence feature alters its chromatin architecture with respect to promoter and intragenic CGIs. We aligned H3K4me1, H3K4me3, H3K27ac and H3K27me3 modifications along reference and alternative promoters and observed different patterns of H3K4me1 and H3K4me3 on HCC patients^6^ (Fig. 4A) and HepG2 cells (Fig. 4B). The CGI promoters have divergent H3K4me3 marks around the nucleosome-depleted region (NDR) at TSSs and flanked by divergent H3K4me1. NonCGI promoters have low levels of H3K4me3 asymmetrically deposited downstream of TSSs, while H3K4me1 marks were non-divergent and enriched at TSSs. This revealed that as the width of H3K4me3 peaks in CG-poor promoters become narrow, the distance between divergent H3K4me1 peaks become shorter and appear continuous. Further, this was strengthened from the CGI viewpoint, as H3K4me3 marks were enriched throughout CGIs and abruptly drop at the end of the CGI (Fig. 4C). In contrast, H3K4me1 peaks were depleted at the CGI body and enriched at their boundaries^36–38^. This pattern is also conserved in enhancers overlapping CGIs (Supplementary Fig. 4A), but different in nonCGI enhancers (Supplementary Fig. 4B). This reinforces that the positional enrichment of H3K4me1 and H3K4me3 is linked to CG-dinucleotides and depletion of H3K4me3 on enhancers is due to majority of enhancers being CG-poor. On the other hand, intragenic CGIs lacked H3K4me3 and H3K27ac modifications (Fig. 4C), thus explaining the absence of transcription. Notably, divergent H3K27ac peaks were enriched on both CG-rich and CG-poor promoters, while they lacked H3K27me3 as expected in active genes (Fig. 4A-B). Further, CGI promoters were enriched for H2A.Z, have a well-defined NDR, and phased +1 downstream the nucleosome, whereas nonCGI promotes have undefined NDR (Fig. 4D). In case of alternative CGI promoters, the gene body histone (H3K79me2, H3K36me3 and H4K20me1) marks presented NDR, while histones were continuous in nonCGI alternative promoters (Supplementary Fig. 4C). As genomic CG dinucleotides influence H3K4me1, H3K4me3, H2A.Z and NDR around TSSs, and as alternative promoters are depleted for CG dinucleotides, their chromatin architecture becomes distinct to that of reference promoters.

**Figure 4.**
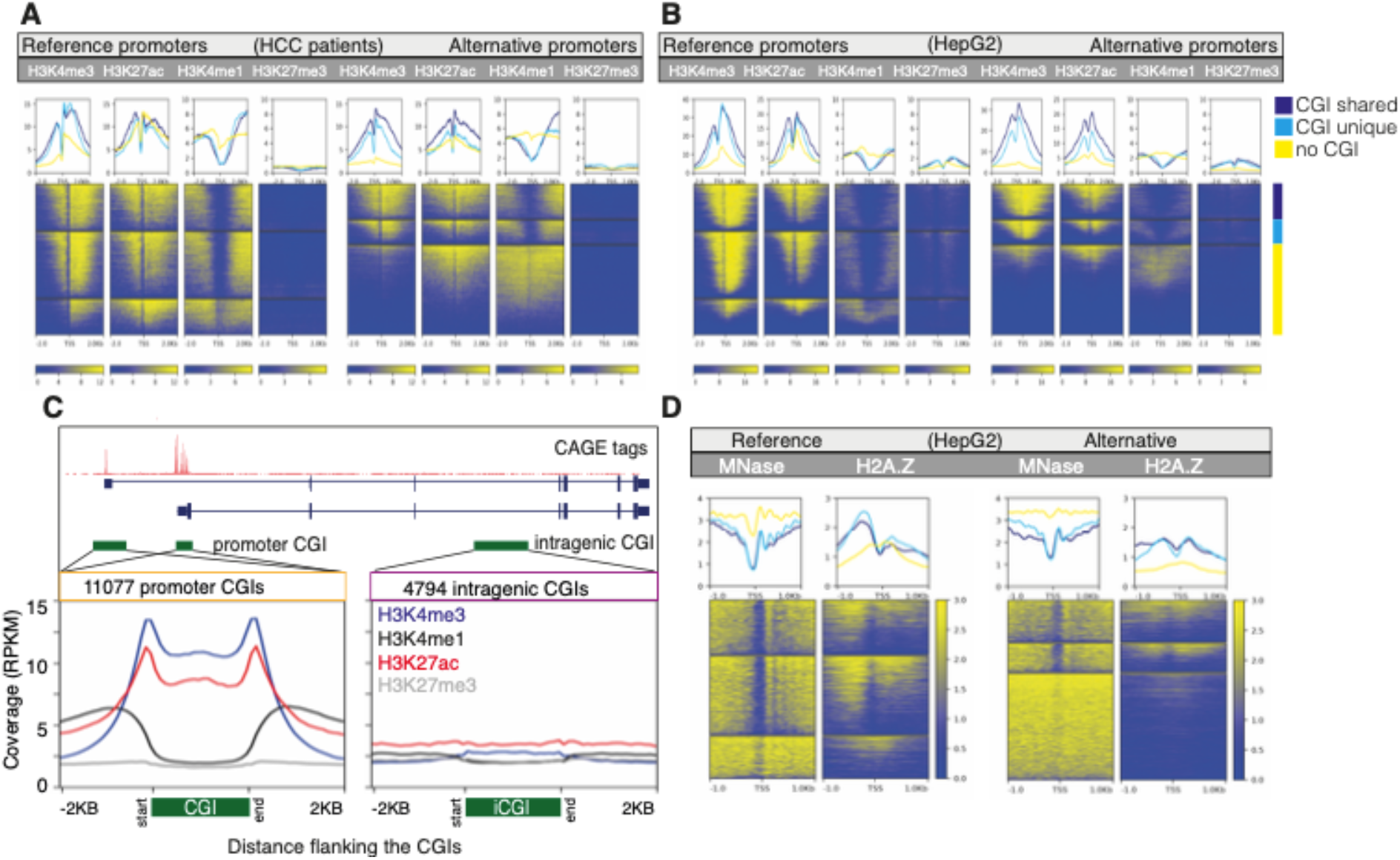
The landscape of histone modifications around reference and alternative promoters. (**A-B**) Line plots show the average histone modification levels of four patients with HCC around reference and alternative promoters. Promoters are classified into three groups based on their overlap with CpG islands (CGIs). CGIs that overlap reference and alternative promoters are classified as shared CGI. Heatmaps show the level of histone modifications for each promoter across shared CGI, unique CGI and nonCGI groups. Representation of data in HepG2 (B). (**C**) A schematic representation of reference and alternative promoters overlapping CGIs along with an intragenic CGI in the gene body. Line plots on the bottom show average coverage of histone marks along promoter CGIs (left) and intragenic CGIs (right). (**D**) Line plots and heatmaps show MNase and H2A.Z levels across reference and alternative promoters on HepG2 cells.

### Dynamic DNA (de)methylation landscapes around alternative promoters

We sought to understand whether DNA methylation facilitates transcription of alternative promoters in CpG-poor regions. The CG dinucleotides are methylated in CpG-poor regions, while CGIs remained unmethylated regardless of the transcriptional state in normal cells^39^ and aberrantly hypermethylated in cancer^2,40^. It is known that H2A.Z marks are mutually antagonistic with DNA methylation^41^. Therefore, as H2A.Z is enriched on CGI promoters (Fig. 4D), these are likewise expected to show a reduced methylation level. Accordingly, reference and alternative promoters overlapping CGIs have low methylation levels at TSSs compared to nonCGI promoters (Fig. 5A). This observation was validated using reduced representation bisulphite sequencing (RRBS) of HepG2 cells (Supplementary Fig. 5A). We next analyzed CG probes (+/− 500 bases around TSSs) and identified hundreds of differentially hypo/hypermethylated promoters (Fig. 5A-B), which significantly overlapped with differentially expressed promoters (Supplementary Fig. 5B). Compared to CGI promoters of expressed genes, aberrant CGI hypermethylation was prevalent among CGI promoters of unexpressed genes (Fig. 5B), suggesting a shift in regulation from H3K27me3 to DNA methylation-based repression^42^. Within CGIs, genomic regions with higher CG density gained methylation, while CGs in non-CGI promoters (in regions with low CG density) lost methylation (Supplementary Fig. 5C). This demonstrates that DNA methylation levels are influenced by the underlying genomic CG dinucleotides at the promoter level.

**Figure 5.**
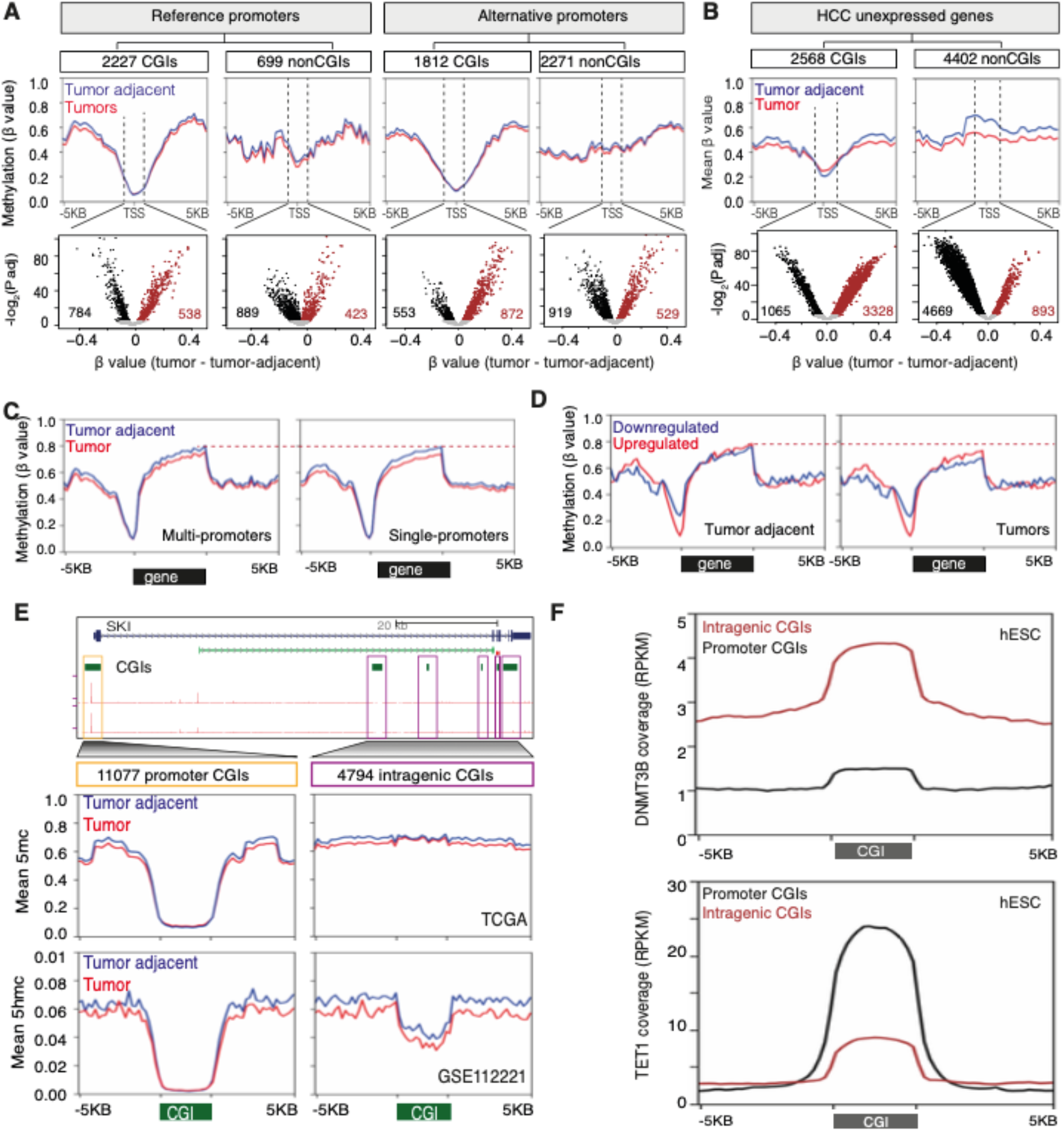
CG dinucleotides influence DNA methylation landscapes differently at reference and alternative promoters. (**A**) Mean methylation levels (β values, top) across the TCGA-LIHC cohort around TSSs of multi-promoter (MP) reference and alternative promoters. Promoters were grouped based on their overlap with CpG islands (CGI). The scatter plot (bottom) shows differentially hypermethylated (brown) and hypomethylated (black) CG probes located 500 bases around TSSs. (**B**) Same as in A, but for unexpressed genes. (**C-D**) Mean methylation levels across TCGA tumors and tumor-adjacent tissues along gene bodies of SP and MP genes (C) and up/downregulated genes. (**F**) A UCSC browser screenshot showing promoter and intragenic CGIs as well as CAGE tags for the *SKI* gene. Zoomed view (top) show the average methylation level of promoter CGIs, intragenic CGIs, and their flanking regions. CGIs of varying lengths are scaled between start and end. Zoomed view (bottom) shows the average demethylation (5hmC) levels. (**E**) Coverage of DNMT3B binding on promoter CGIs and intragenic CGIs across human ES cells. (**F**) Coverage of TET1 binding on promoter and intragenic CGIs across human ES cells.

Gene bodies are known to have high levels of DNA methylation^20,43^. However, it is unclear in what way the intragenic CG density influences DNA methylation in tumors that preferentially facilitates transcription in low CG regions. Intragenic regions have high methylation levels across all promoters, but globally this was decreased in HCC compared to the tumor-adjacent tissues (Fig. 5C). Decreased intragenic methylation was more evident among downregulated promoters (Fig. 5D) and most notably among genes without intragenic CGIs (Supplementary Fig. 5D), suggesting that the local CG density influences intragenic (de)methylation in tumors. In fact, intragenic CGIs have high methylation, which is opposite to promoter CGIs in HCC patients (Fig. 5E) and in HepG2 cells (Supplementary Fig. 5E). Compared to CG-poor flanking regions, the methylation level of intragenic CGIs remained globally unchanged in HCC tumors (Fig. 5F), which is due to high levels of bound DNMT3B (Fig. 5F). Intragenic CGIs have low levels of DNA demethylase (5hmC levels) activity (Fig. 5E) measured by TET-assisted reduced representation bisulphite sequencing (TAB-RRBS)^6^, which is consistent with low levels of TET1 binding (Fig. 5G). Thus, the lack of active demethylation on intragenic CGIs can explain its sustained high level of methylation to maintain the repressive state, resulting in lack of activation of alternative promoters from intragenic CGIs (Fig. 4D). In contrast, alternative promoters with CG-poor regions are sensitive to demethylation in tumors, which in turn may facilitate an open chromatin structure and transcription initiation.

### Dysregulation of SETD2, H3K36me3 and DNA methylation facilitates pervasive open chromatin and subsequent Pol2 binding

The methyltransferase SETD2 deposits H3K36me3^44^ after Pol2 passage^45^, which in turn recruits DNMT3B^21^ to maintain a repressive chromatin state^46^. Thus, we reasoned that dysregulation of *SETD2* may alter the repressive chromatin state and facilitates alternative transcription. Old yeast cells have decreased H3K36me3 levels and increased intragenic transcripts^47^, while DNA methylation decreases with age in humans^48^. HCC tumors showed reduced DNA methylation within gene bodies (Fig. 5C). H3K36me3-modified nucleosomes were enriched along gene bodies and globally reduced by *SETD2* knockdown (Fig. 6A). Within gene bodies, CG-rich regions have higher H3K36me3 marks also in *SETD2* knockdown (Fig. 6B). In HepG2, we observed lower H3K36me3 marks compared to in normal hepatocytes (HepaRG) (Supplementary Fig. 6A, see methods). As H3K36me3 recruits DNMT3B^21^, the gene body coverage of H3K36me3 (Supplementary Fig. 6B) coincides with DNMT3B coverage (Fig. 6C). Using CRISPR epitope tagging (through insertion) of DNMT3B^49^ revealed DNMT3B signals decreased along the gene body (Supplementary Fig. 5C) and showed a pattern of repressive epigenetic marks along the gene body.

**Figure 6.**
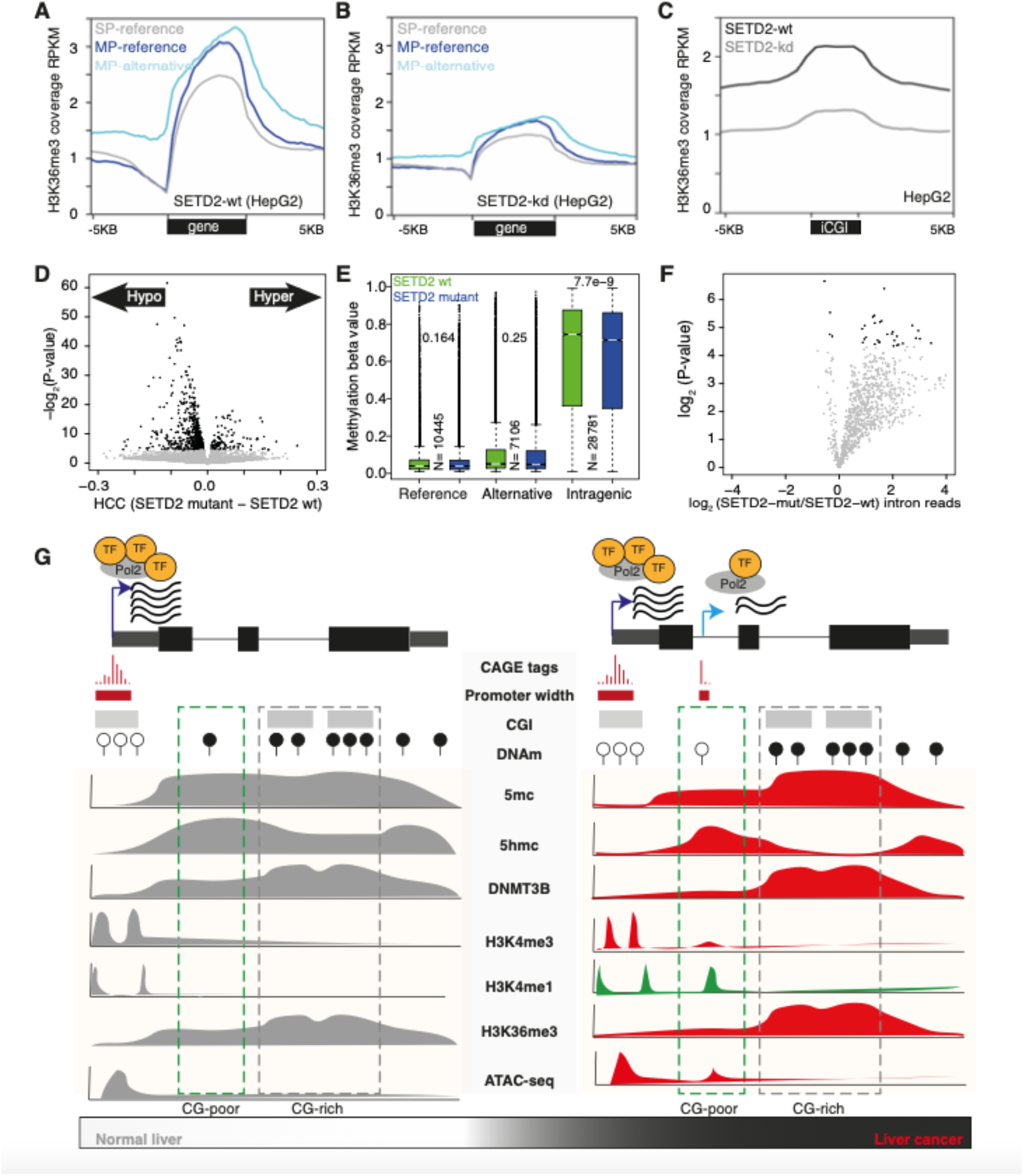
Regulation of SETD2. (**A-B**) The average coverage of H3K36me3 along gene body and flanking regions of single promoter (SP) and multi-promoter (MP) genes in SETD2-wt and SETD2-kd in HepG2 cells. (**C**) The average coverage of H3K36me3 along intragenic CGIs and flanking regions in SETD2-wt (A) and SETD2-kd (B) in HepG2 cells. (**D**) Volcano plot shows hypermethylated and hypomethylated CG probes between SETD2-mutant and SETD2-wt tumors from TCGA-LIHC. (**E**) Boxplot shows average DNA methylation levels of CG probes around reference and alternative promoters and intragenic regions. (**F**) Volcano plot shows fold-change of intronic reads in SETD2-mutant versus SETD2-wt. (**G**) Schematic diagram to show transcription of alternative promoters in HCC. The chromatin structure of intragenic CG-rich and CG-poor regions have different distribution of 5mC, 5hmC, H3K36me3 and DNMT3B, leading to pervasive initiation of the alternative promoter from CG-poor regions in HCCs.

We reasoned that if *SETD2* is deregulated in patients, this would lower H3K36me3 levels, which in turn would lower DNMT3B binding and thus, decrease in DNA methylation across gene bodies (Fig. 5D). To support our hypothesis, we asked whether *SETD2* mutations accelerated DNA demethylation in HCC patients. Compared to *SETD2* wildtype tumors (n=352), *SETD2* mutated tumors (n=16) were significantly hypomethylated (Fig. 6D and Supplementary Fig. 6D) which were enriched in gene bodies (Fig. 6E) compared to promoters. Notably, SETD2-mutants have a high level of intronic reads (Fig. 6F), providing evidence for transcribed reads from introns. Predominant hypomethylation observed across gene bodies in tumors was more prevalent in SETD2-mutants, as *SETD2* mutations accelerate the epigenetic control, compromising the maintenance of DNA methylation, and result in pervasive activation of alternative promoters. Thus, the interplay of DNA methylation, chromatin accessibility and histone modification in cancer preferentially facilitates alternative transcription in CG-poor regions (Fig. 6G).

## Discussion

We have comprehensively annotated 4083 (3052 annotated and 1031 novel) alternative promoters in HCC using high-resolution CAGE-seq, providing a framework for functional studies. These alternative promoters were supported by multiple lines of evidence from orthogonal sequencing assays across different HCC patient cohorts and in HepG2 cells. Currently, there is no consensus on which promoter is assigned as reference or alternative, though a highly expressed promoter is often annotated as the reference promoter^16,19^, which is what we used in our analysis. In addition, we ensured that the reference promoter was expressed in normal livers. Around 40% of alternative promoters were unexpressed in normal liver and hence will remain as alternative promoters. Thus, only a small fraction of alternative promoters that have comparable expression levels like the reference promoters can vary as either reference or alternative promoters across cohorts.

We found that the CG dinucleotide density is the major distinguishing feature between reference (has a high CG density) and alternative (has a low CG density) promoters. Human promoters are mostly CG-rich^32^ and have distinct promoter architecture and regulation^11^ compared to tissue-specific CG-poor promoters^17^, which we validated pan-cancer. This highlighted the unexpected presence of two distinct regulatory promoter architectures within a gene, showing it is widespread in cancer. Most alternative promoters are CG-poor and detected in lower proportion across other tumor types, indicating they drive tissue-specific or cell type-specific transcription. Their tissue-specific expression was further supported by the enrichment of liver-specific transcription factors (for example HNF4A). While reference promoters are generally ubiquitous and represent the predominant degree of transcription, it is important to understand the cell type transcriptional contribution that drive tissue-specific alternative promoters. This requires implementation of single cell capturing the 5’ends^50^, while single cell methods currently in use are enriched towards genes at the 3’-ends and therefore, insensitive for promoter detection. Single cell capturing 5’-ends in mouse neurons^50^ have revealed many alternative promoters that were detected in bulk tissues, which suggest that most alternative promoters might be co-expressed in the same cell.

The epigenetic (histone modifications and DNA methylation) landscapes of reference and alternative promoters are dramatically different, which is primarily due to their differences in CG dinucleotide frequency. The H3K4me3 marks are enriched on CGIs^51^, while we showed that intragenic CGIs lack H3K4me3 marks due to retention of high DNA methylation in the tumor. It is known that aged cells (and cancer cells) generally loose DNA methylation from gene bodies^48^. We described a hierarchy that explained preferential loss or retention of intragenic DNA methylation dependent upon the flanking CG dinucleotide density where CG-rich regions retain high tumor DNA methylation. This preferential retention of intragenic DNA methylation is regulated by *SETD2*, H3K36me3 and DNMT3B as they preferentially bind to CG-rich regions. This leads intragenic CG-rich regions with more repressive chromatin marks and that would require higher levels of DNA demethylating enzyme to make their chromatin accessible. Instead, these intragenic CG-rich regions have low levels of *TET1* binding and thus, have the reduced active demethylation. Thus, intragenic CGIs are less favorable to act as alternative promoters in cancer, though they can act as alternative promoter during embryonic development and cell differentiation^33^. In contrary, intragenic CG-poor regions have lower levels of repressive chromatin state and higher levels of *TET1* binding, which collectively makes their DNA sensitive to active demethylation and accessible chromatin to experience pervasive Pol2 binding^52^ that upon availability of a tissue-specific transcription factor will efficiently initiate and elongate these Pol2. We propose that other tumors have similar mechanism in activating alternative promoters, while their preferred location will be dependent upon DNA demethylated sites that are enriched for tissue-specific transcription factors of the specific tumor tissue.

The general view on why some genes have multiple promoters is that a cell needs more expression of specific genes and transcription from multiple promoters may add flexibility manifested through different molecular mechanisms in regulating gene expression^53^. This view is mostly based on our understanding of alternative promoters’ usage in normal cells. However, in cancer, alternative promoters are pervasively activated, raising an interesting question as to whether it is advantageous for tumor cells. We propose different scenarios where alternative promoters offer functional advantage needed for the tumor. Firstly, alternative promoters can downregulate reference promoters through transcriptional interference, which often coincide with important genes including hepatocyte markers (Supplementary Fig. 2F). Accelerated loss of expression levels of hepatocytes markers help hepatic cells to lose their cellular identify and gain new cellular identify as the malignant tumor. To develop the cellular identity, tumor cells need to utilize signaling pathways differently in a globally changed cancer background where the alternative promoters provide better adaptation for the cancer regulatory systems driven by a distinct promoter architecture. As such, alternative promoters detected only in tumors were enriched for genes in signaling pathways (Fig. 2C), which globally did not downregulate reference promoters (Fig. S2F). This suggests that signaling pathway genes co-opt both promoters, where tumor cells utilize signaling pathways differently via alternative promoter usage (for example in *CDKN2A* (Supplementary Fig. 1H) and *ERBB2* (Supplementary Fig. 1I)). On the clinical perspective, MP genes are associated with shorter survival time and alternative promoters were better in predicting overall survival. This opens an avenue to elucidate improved diagnostic biomarkers for early onset of cancer as many alternative promoters are expressed in a tumor-specific manner. Moreover, signaling pathways genes are often potential targets in designing drug(s), hence we should focus on target tumor-specific alternative transcripts rather than the reference transcript. While our analyses suggest an important role of alternative promoters in liver cancer biology, their biological significance in the full spectrum of liver cancer remains unexplored. The selective use of alternative promoters often goes uncharacterized in gene expression analyses in standard RNA-seq analyses. We strongly encourage scientists in the liver community to carefully inspect whether their genes of interest are under alternative promoter regulation and whether tumor phenotype of a gene is primarily driven by alternative promoter.

## Methods

### Mapping of HCC CAGE-seq reads to define CAGE transcription start sites (CTSSs)

Raw CAGE-seq reads were downloaded from a previous study^8^. The CAGE-seq reads were mapped to the human genome (hg19) with bowtie2^54^ by allowing up to two mismatches. On average, around 80-90% of sequenced reads were mapped, resulting in an average of 15-16 million mapped reads (Supplementary Table 1). The 5’ end of mapped CAGE reads provide transcription start sites (TSSs) at single nucleotide resolution and is termed as CAGE transcription start sites (CTSSs). Low quality and multi mapping CTSSs with MAPQ score below 20 were filtered using SAMtools^55^. The CTSSs that overlapped with 421 blacklisted regions from ENCODE^56^ were excluded. The CAGE protocol often adds “G” at the 5’ end of capped TSS^11^, which generally remain unmapped and shift transcription start site by 1 nucleotide. We used SAMtools and detected mapped reads with unmapped “G” at the first base. We then corrected such CTSSs by shifting the TSS position by 1 nucleotide as defined before^12^.

### Transcript clusters (TCs) and generating consensus TCs across cohort

To define transcript clusters (TCs) for each patient, we clustered CTSSs in same strand that overlapped within 20 nucleotides^12^. The CTSSs with the highest expression level within TC was defined as the dominant CTSS. All CTSSs within the TC were added that defined the expression level of TC. We then computed the interquartile width of the TCs by trimming the edges of the TCS in the range of 0.1 to 0.9 percentile of the TC. To define the consensus TCs across the cohort, we clustered TCs from all patients and computed their expression levels. To ensure that consensus TCs have robust expression levels, we retained TCs only of their expression was higher than 1TPM in at least 15 samples (15% of cohort) and had a minimum of 3 TPM at least in 1 sample.

#### Assignment of reference and alternative promoters among annotated transcripts

A gene with a single promoter was classified as a single promoter gene and its promoter is assigned the reference (alias as primary, major, main) promoter. Genes with two or more promoters were classified as multi-promoter genes. The promoter with highest mean expression level at the population level that is also expressed in normal liver tissue was assigned as reference promoter. The remaining promoters of that gene was assigned as an alternative (alias minor) promoters of that gene. The terms reference TSS and alternative TSS have been used ambiguously.

#### Annotation of novel alternative promoters

To annotate novel alternative promoters, we implemented a pipeline (**Fig. 1C**) that systematically filtered intragenic TCs that are unlikely to represent true promoters. We filtered CAGE TCs overlapping small RNAs, drosha processing site of pre-miRNAs^23^ and 5’ ends capping of snoRNAs^17^. Thousands of intragenic CAGE TCs are detected within exons which represent post-transcriptional processing and characterized by GG-initiation^11,12^. We filtered intragenic CAGE TCs with GG-initiation and those overlapping with coding exons. We filtered intragenic CAGE TCs that overlapped with annotated enhancers^24,57^. We also annotated novel enhancers from CAGE-seq by computing directionality score for each TCs as described previously^24^. Briefly, directionality score (DS) for intragenic TCs were calculated by measuring expression level of TCs in forward and reverse strand, where DS=(Forward-Reverse)/ (Forward+Reverse). CAGE TCs with directionality score between 0.5 and −0.5 were classified as enhancer RNAs. On the remaining TCs, we only retained those that have YR-initiation or YC-initiation initiation motif^15^. The remaining 1077 TCs represent high-confidence true promoter tags that act as alternative TSSs to annotated genes. We clustered proximal TCs with 300 bases thus resulting in 1031 novel alternative TSSs. To provide evidence of RNA transcripts for these 1031 alternative TSSs, we analyzed transcript models from RNA-seq and expressed sequence tags from UCSC database^58^.

### Mapping and visualization of histone ChIP-seq of HCC patients and HepG2 cells

We downloaded the raw sequence reads of H3K4me1, H3K4me3, H3K27ac and H3K27me3 ChIP-seq data for four HCC patients^6^. We downloaded raw sequence reads of H3K36me3 marks on SETD2 wild type and SETD2 knockdown conditions on HepG2 cells^59^. The raw sequence reads were mapped using bowtie2^54^ and excluded multi mapping reads. We used deepTools^60^ to generate coverage tracks which are normalized as RPKM.

### Analysis of DNA methylation of HCC patients and HepG2 cells

The Illumina 450K DNA methylation data from TCGA HCC cohort was downloaded from UCSC Xena hub^61^. The methylation levels were computed as beta value in the range of 0-1. The reduced representation bisulphite sequencing (RRBS) of HepG2 cells were downloaded from ENCODE. The TET-assisted reduced representation bisulphite sequencing (TAB-RRBS) of HCC patients were downloaded as beta value^6^. For each CG probes, we computed the average methylation beta value across the cohort, separately for tumor and tumor-adjacent tissues. To identify differentially methylated CG probes around promoter regions, we performed t test on individual methylation levels between tumor and tumor adjacent tissues. The p-value were adjusted for multiple correction and p-value less than 0.05 was defined as significantly methylated CD probe. For visualization of average methylation levels around TSSs and CGIs, the mean beta value was converted into bigwig tracks and plotted average beta value using deepTools^60^.

### Enrichment of transcription factor and *de-novo* motif analysis

We downloaded transcription factors (TFs) ChIP-seq peaks for HepG2 from ENCODE^62^. To calculate the density of TFs, we intersected TFs peaks with promoter regions by using bedtools^63^. Different TFs overlapping promoters were summed to determine the total number of TFs per promoter.

To identify overrepresented transcription factors, we performed motif analyses using HOMER^64^. We used 500 bases around transcription start sites as the search region to detect motifs. The background regions were controlled for nucleotide composition and selected by default by HOMER. We first identified motifs using the following parameter “findMotifsGenome.pl -chopify -len 8,10,12 -S 25 - size −500,500”. To compare these identified motifs with known motifs, we reran prediction using the following command “findMotifsGenome.pl -chopify -len 8,10,12 -S 25 -mcheck -mknown -size - 500,500”. For visualization of motifs, we computed the matrix of p-value across promoter types and plotted them as heatmaps.

### Signature genes from the literatures

Curated signature genes for hepatocytes^25^, Hoshida signature genes^26^, sorafenib resistance signature^28^ and GSEA cancer hallmark genes^27^ were downloaded from literatures. We interested these gene signatures with annotated single-promoter (SP) and multi-promoter (MP) genes. We computed Fisher’s exact test to determine the statistical significance of overlap between signatures genes with single and multi-promoter genes.

### Patients’ survival analysis

To compute the overall survival of patients associated with expression level for each transcript, we downloaded TCGA LIHC clinical data^2,65^ and transcript expression levels from UCSC Xena hub^61^. Each transcript was sorted based on their expression levels and classified into high-expression and low-expression groups. We associated the expression levels of these transcripts from two (high-expression and low-expression) groups with survival status of patients and performed Kaplan-Meier analyses. For multi-promoter genes, we performed Kaplan-Meier analysis on reference transcript and on alternative transcript.

### Differential expression at promoter and gene level

Differentially expressed genes and promoters were identified using DE-seq2^29^. The significance cut-off was defined at adjusted p-value of 0.05 and fold-change of 2. For gene level analysis, we summed the expression levels of reference and alternatives promoter for each multi-promoter genes.

### Analysis of N-terminus of protein and UniProt domains

To compare whether alternative promoter altered the N-terminus of protein, we analyzed only those alternative transcripts that have assigned UniProt protein domains, and hence excluded noncoding transcripts and novel alternative promoters. We compared the N-terminus of the reference and alternative promoters, and if they had different start codon, they were assigned as different N-terminus protein.

## Data availability

All sequencing data analyzed in this study are publicly available from previous studies. The results are in part supported by data generated by the TCGA Research Network: https://www.cancer.gov/tcga, FANTOM5, ENCODE and NIH Roadmap Epigenome. Data accession codes are provided. All processed data are provided as supplementary table and UCSC data tracks.

## Acknowledgements

We thank Prof. Ferenc Mueller, Dr. R. Taylor Raborn, Dr. Colm J O’Rourke and Prof. Cord Brakebusch for critical reading of the manuscript. We thank Dr. Sachin Pundhir, Dr. Monika Lewinska, Dr. Juan F Lafuente Barquero and Dr. Yuewan Luo for critical discussion and suggestions.

---

**Supplementary Figure 1.**
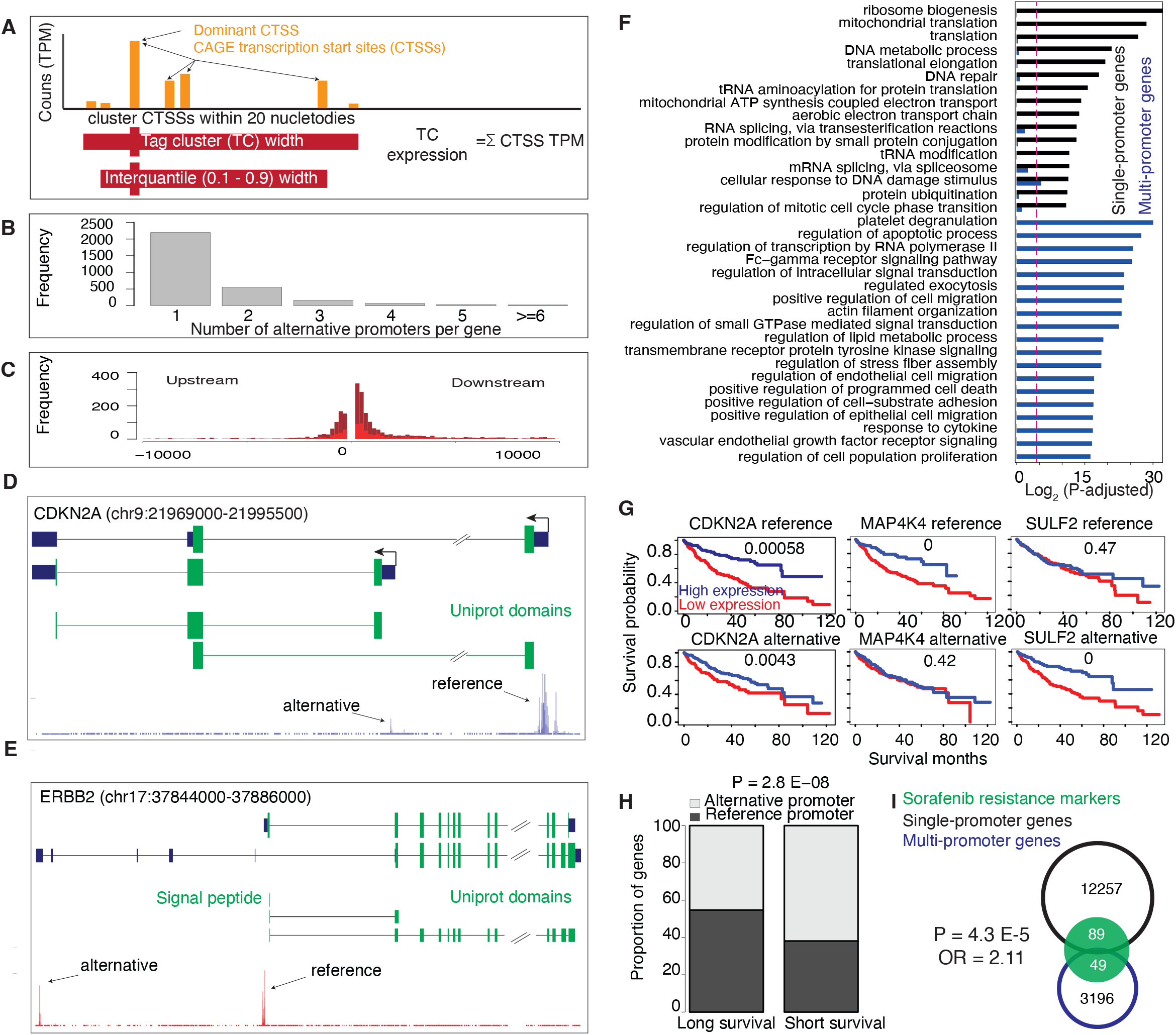
Annotation of alternative promoters in patients with hepatocellular carcinoma (HCC). (A) A schematic workflow that describes the clustering of proximal CAGE transcription start sites (CTSSs) into transcript clusters (TCs). The height of CTSSs determines the frequency of mapped reads and is used to quantify the expression of CTSS in tags per million (TPM). The expression of TCs is determined by the sum of all overlapping CTSSs. The interquartile widths of TCs are determined by 0.1 to 0.9 fraction of expression levels. (B) Barplots showing the number of alternative promoters for each multi-promoter genes. (C) Barplot showing the distance between reference and alternative promoters. (D-E) A UCSC browser screenshot of CDKN2A (D) and ERBB2 (E) genes along with tracks of uniprot domains and CAGE-seq. Uniprot domains show alternative promoters miss a segment of protein domains. (F) Enriched gene ontology terms associated with single-promoter and multi-promoter genes. (G) Individual examples of Kaplan-Meier survival analysis for expression of reference and alternative promoters. (H) Classification of survival-associated genes based on time to death (long/short survival time) for reference and alternative promoters. Alternative promoters are significantly (P= 2.8E-08; Fisher’s exact test) associated with short survival. (I) Overlap of sorafenib resistance marker-genes with single and multi-promoter genes revealed an enrichment (P= 4.3E-05; Fisher’s exact test) of marker-genes among multi-promoter genes.

**Supplementary Figure 2.**
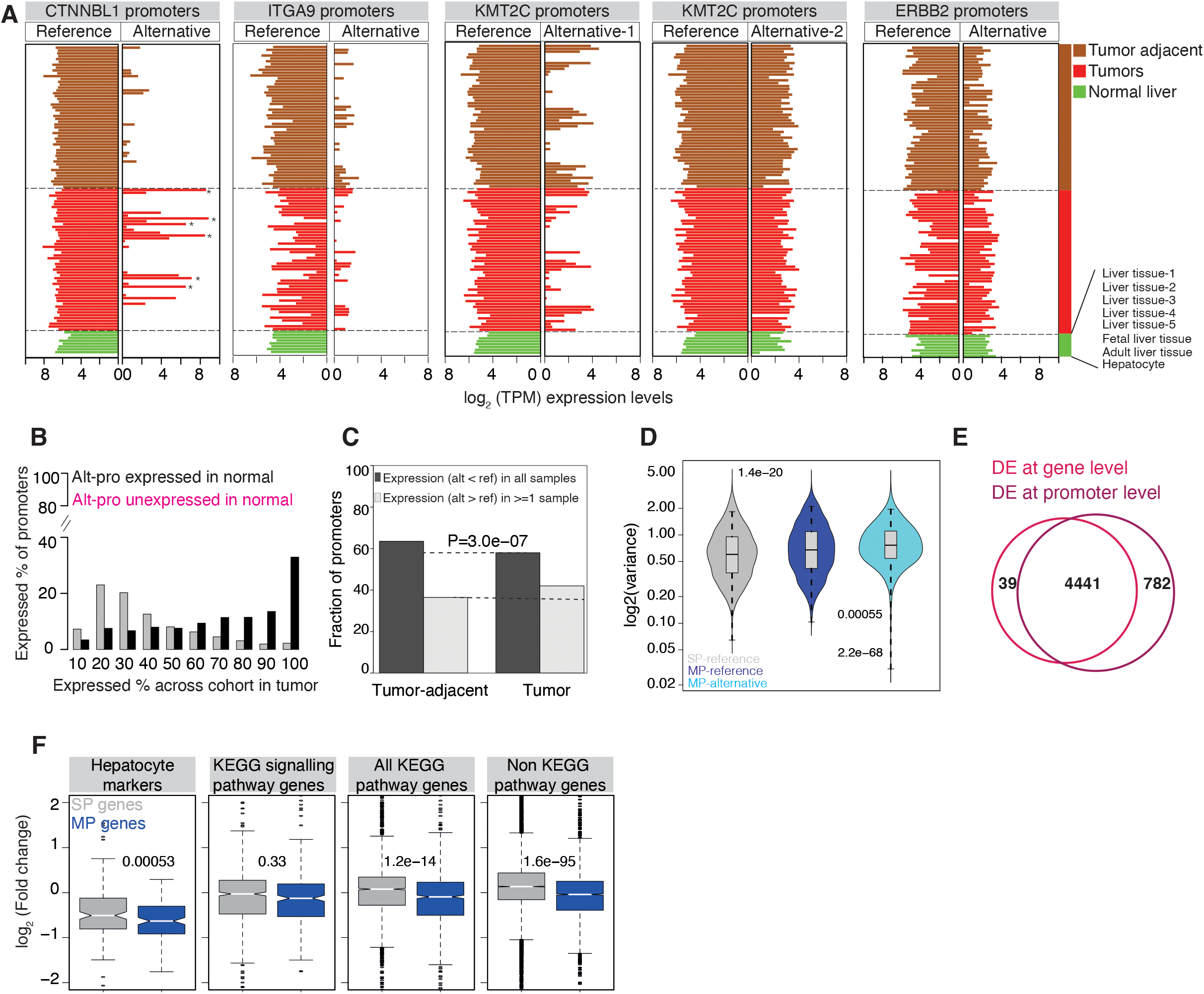
Expression dynamics of alternative promoters. (A) Barplots show the expression level of reference and alternative promoters for selected genes (CTNNBL1, ITGA9, KMT2C and ERBB2). Two plots of the gene KMT2C represent two alternative promoters. Each bar represents an individual sample within the data set including 50 HCCs, matched tumor-adjacent tissues and 8 normal liver tissue samples. Each sample is ordered in the same manner across all plots. The asterisk indicates when the expression level of an alternative promoter is higher than its reference promoter within a sample. (B) Distribution of alternative promoters expressed across the cohort. Alternative promoters were classified into two groups based on the presence or absence of their expression in the normal liver tissues. X-axis indicates the percentage a promoter is expressed across the cohort. (C) Barplots show the distribution of expression levels of reference and alternative promoters at individual samples. (D) Barplot show the distribution of variance of expression levels across the cohort for both reference and alternative promoters. (E) Venn diagram represents the overlap of differentially expressed genes and promoters between tumor and tumor-adjacent tissues. (F) Expression fold change between tumor and tumor adjacent tissues for single-promoter (SP) and multi-promoter (MP) genes across different functional classification of genes. P values were computed using the t

**Supplementary Figure 3.**
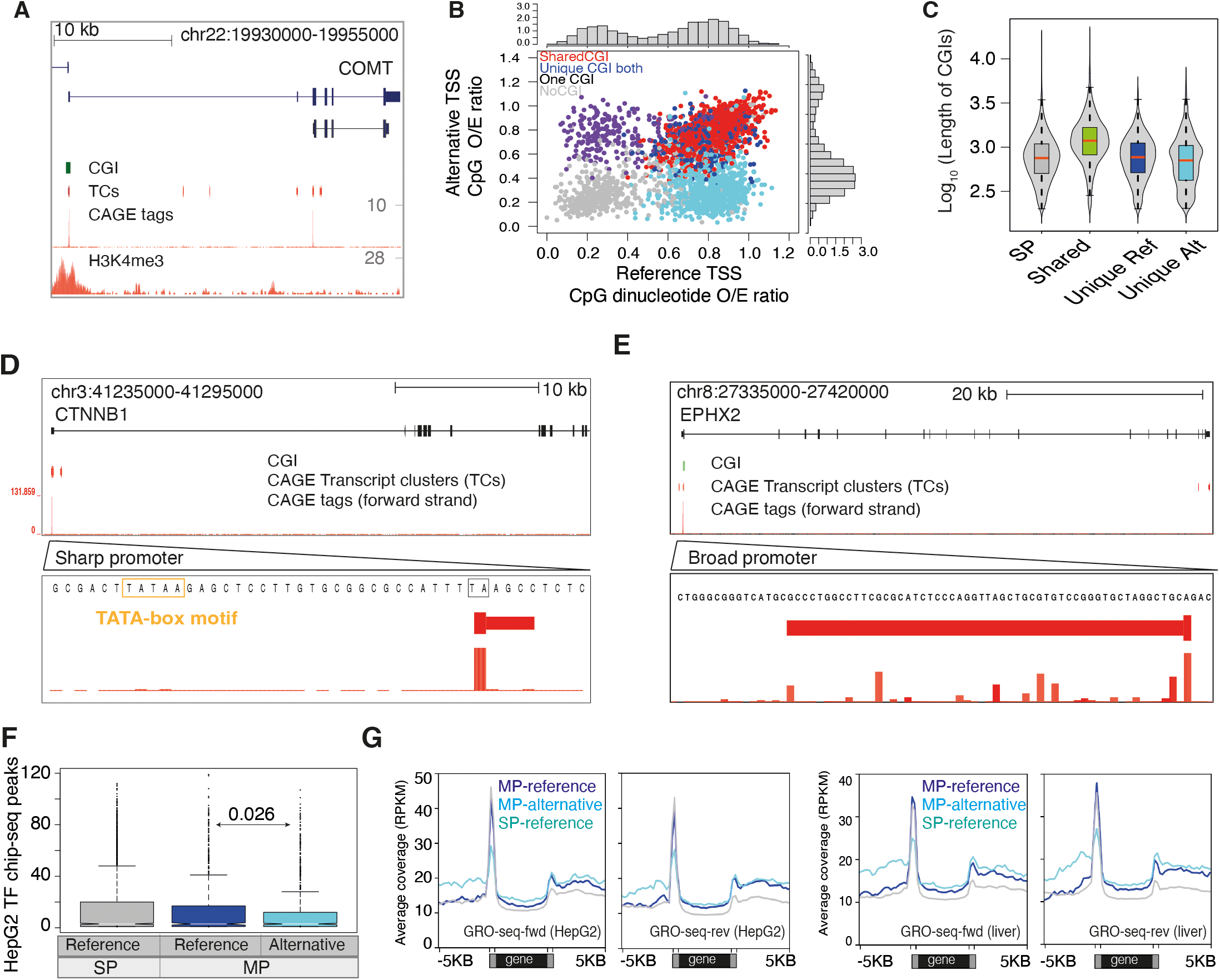
Sequence and motif enrichment of reference and alternative promoters. (A) A UCSC browser screenshot of COMT gene along with tracks of CpG islands (CGI), CAGE-seq and H3K4me3. (B) Scatter plot showing the observed/expected (O/E) ratio of CG dinucleotides around reference and alternative promoters. (C) A distribution of CGI lengths at overlapping promoters. CGIs shared by reference and alternative promoters were significantly longer. (D-E) A UCSC browser shot of CTNNB1 and EPHX2 genes along with tracks of CAGE tags, transcript clusters and CGI. The zoomed view shows the width of transcript clusters and sequences around dominant transcription start site (TSS). The CTTNB1 gene has sharp promoter with only one dominant TSS and TATA box motif in upstream region. The EPHX2 gene has broad promoter with multiple TSSs and lack TATA box motif in upstream region. (F) Frequency of ENCODE transcription factors peaks on reference and alternative promoters of single promoter (SP) and multi-promoter (MP) genes. (G) The average coverage of nascent RNA reads from GRO-seq in HepG2 cells for genes in forward and reverse strand.

**Supplementary Figure 4.**
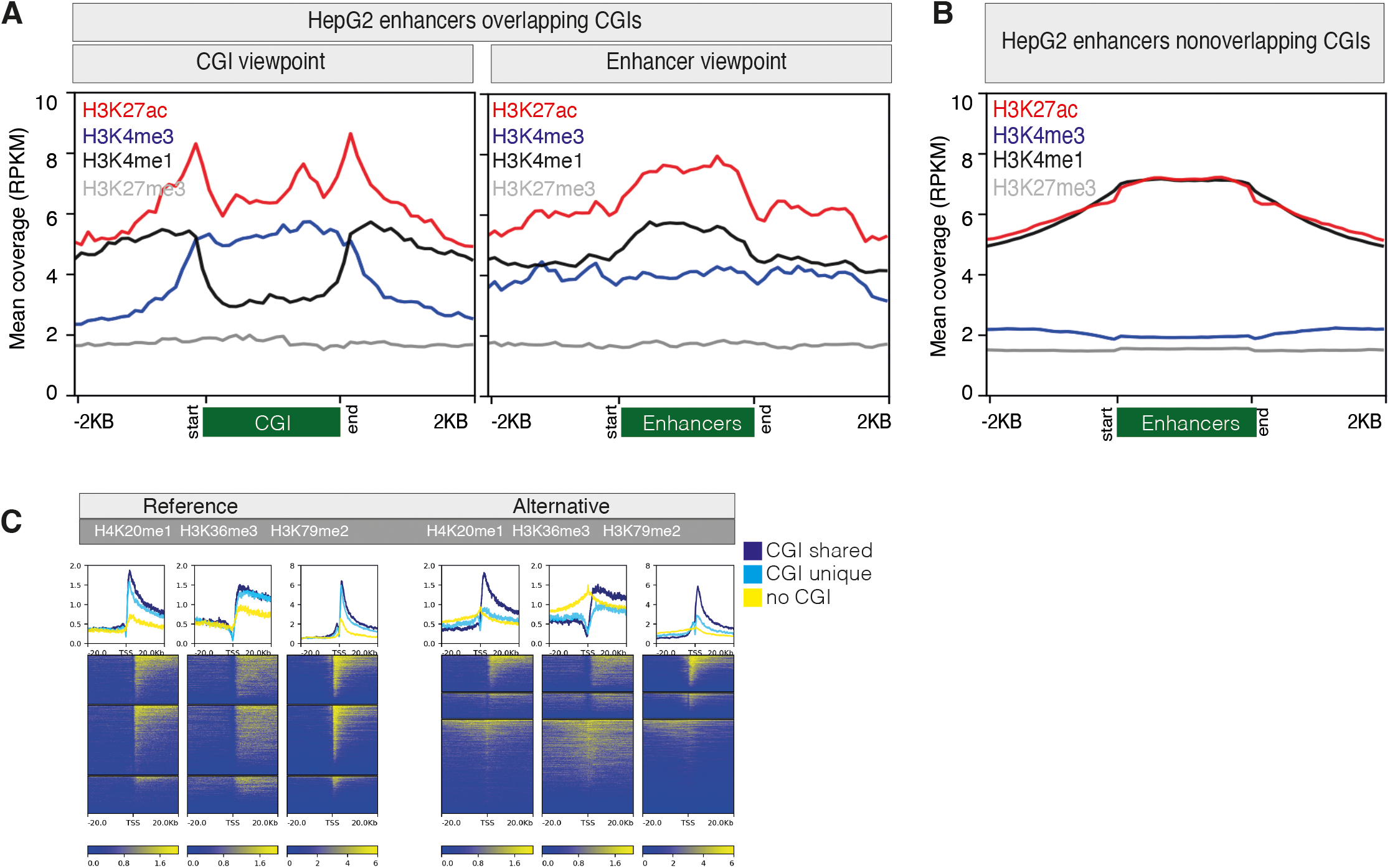
Histone modifications around CGI enhancers, nonCGI enhancers along with reference and alternative promoters. (**A**) Average coverage and positional enrichment of four histone modifications (H3K4me1, H3K4me3, H3K27ac, H3K27me3) on HepG2 enhancers overlapping CGIs. Left panel shows histone coverage along the start and end of CGIs overlapping enhancers. Right panel shows histone coverage along the start and end of enhancers within CGIs. (**B**) Average coverage and positional enrichment of four histone modifications (H3K4me1, H3K4me3, H3K27ac, H3K27me3) along nonCGI enhancers. (**C**) Average coverage of histones (H4K20me1, H3K26me3 and H3K79me2) along reference and alternative promoters.

**Supplementary Figure 5.**
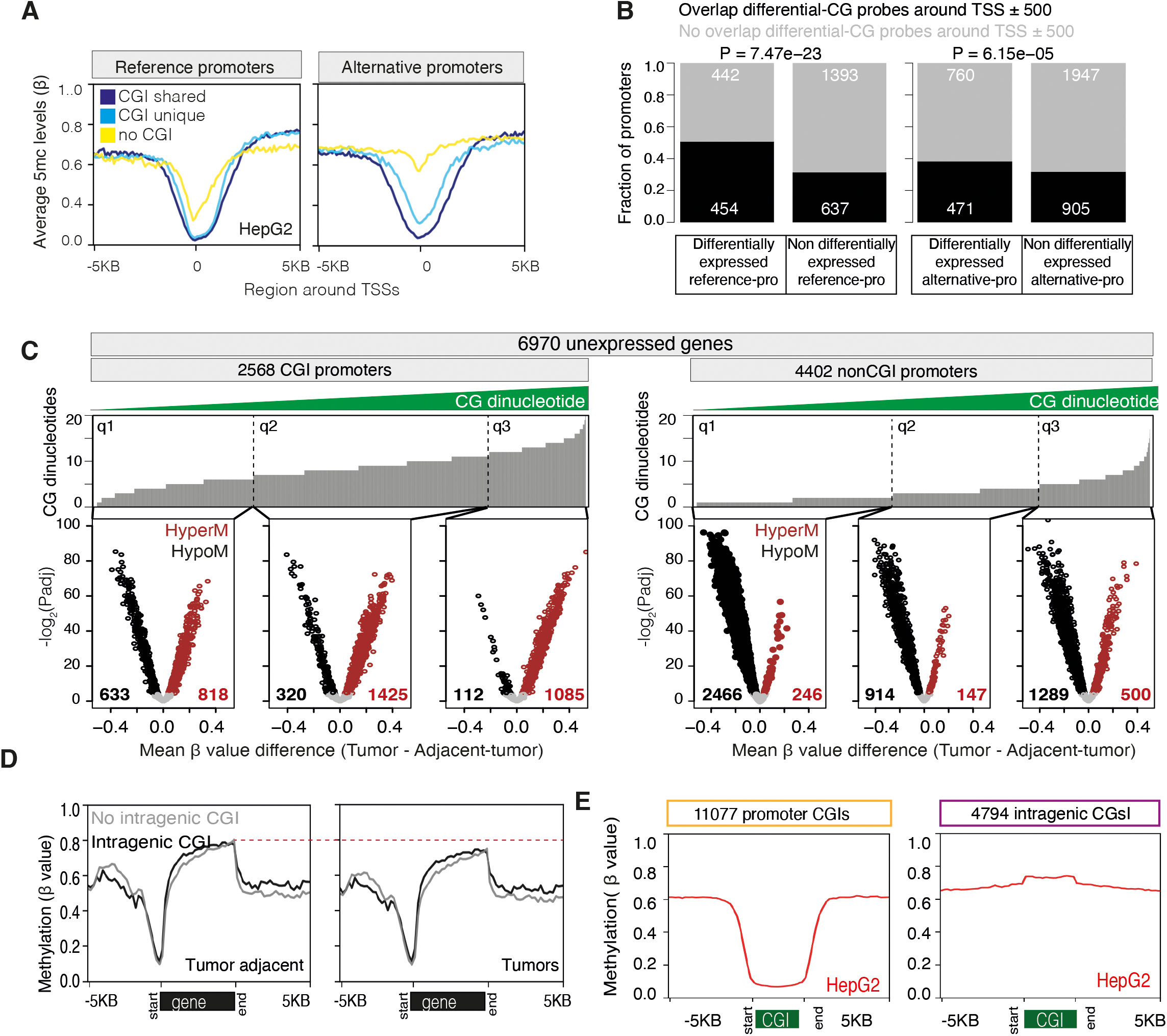
The DNA methylation landscape around reference and alternative promoters. (A) Line plots show the methylation level (β values) obtained from reduced representation bisulfite sequencing (RRBS) on HepG2 cells across reference and alternative promoters, which are grouped based on their overlap with CpG islands (CGIs). (B) Overlap of differentially expressed reference and alternative promoters with differentially methylated (DM) CG probes in a 500 base pair window around TSSs. (C) DNA methylation profiles across TCGA-LIHC among unexpressed genes classified into two groups based on their overlap with CGIs. Barplots (top) show the frequency of CG dinucleotides in a 100-nucleotide window for each CG probe in promoter regions of unexpressed genes. CG probes are ordered based on increasing number of CG dinucleotides in 100 nucleotide windows and divided into three quartiles (q1, q2, q3). The scatter plots (bottom) show differentially hypermethylated (brown) and hypomethylated (black) CG probes in 500 bases around TSSs. (D) Mean methylation level of promoter and intragenic CGIs across HepG2 cells. (E) Mean methylation level along promoter CGIs and intragenic GIs on HepG2 cells.

**Supplementary Figure 6.**
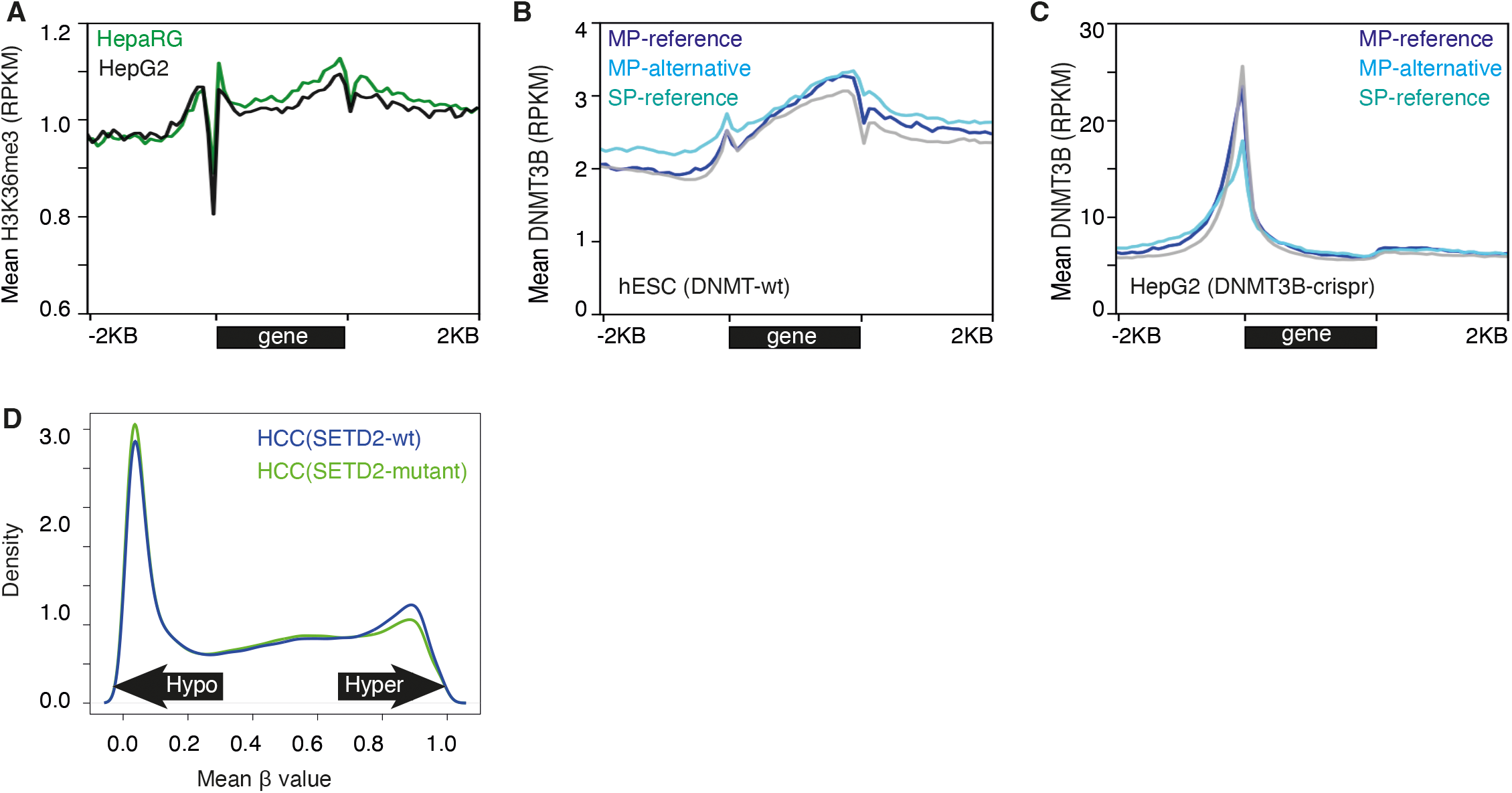
Genomic coverage of H3K36me3, DNMT3B and RNA Pol2 across the gene body. (A) The average coverage of H3K36me3 along the gene body and flanking regions in HepG2 cells and HepaRG cells (normal hepatocytes). (B) The average coverage of DNMT3B across the gene body in human embryonic stem cells (hESC). (C) The average coverage of DNMT3B across the gene body in CRISPR epitope tagging (through insertion) of DNMT3B in HepG2 cells. (D) The density plot shows the distribution of mean methylation levels (β values) of HCC patients (n=352) with SETD2 mutations and HCC patients (n=16) with wildtype SETD2.

## References

1. Sung, H. et al. Global Cancer Statistics 2020: GLOBOCAN Estimates of Incidence and Mortality Worldwide for 36 Cancers in 185 Countries. CA Cancer J Clin 71, 209–249 (2021).

2. Cancer Genome Atlas Research Network. Electronic address, w.b.e. & Cancer Genome Atlas Research, N. Comprehensive and Integrative Genomic Characterization of Hepatocellular Carcinoma. Cell 169, 1327–1341 e23 (2017).

3. Villanueva, A. et al. DNA methylation-based prognosis and epidrivers in hepatocellular carcinoma. Hepatology 61, 1945–56 (2015).

4. Schulze, K. et al. Exome sequencing of hepatocellular carcinomas identifies new mutational signatures and potential therapeutic targets. Nat Genet 47, 505–511 (2015).

5. Fujimoto, A. et al. Whole-genome mutational landscape and characterization of noncoding and structural mutations in liver cancer. Nat Genet 48, 500–9 (2016).

6. Hlady, R.A. et al. Integrating the Epigenome to Identify Drivers of Hepatocellular Carcinoma. Hepatology 69, 639–652 (2019).

7. Corces, M.R. et al. The chromatin accessibility landscape of primary human cancers. Science 362(2018).

8. Hashimoto, K. et al. CAGE profiling of ncRNAs in hepatocellular carcinoma reveals widespread activation of retroviral LTR promoters in virus-induced tumors. Genome Res 25, 1812–24 (2015).

9. Lin, Z. et al. RNA m(6) A methylation regulates sorafenib resistance in liver cancer through FOXO3-mediated autophagy. EMBO J 39, e103181 (2020).

10. Lenhard, B., Sandelin, A. & Carninci, P. Metazoan promoters: emerging characteristics and insights into transcriptional regulation. Nat Rev Genet 13, 233–45 (2012).

11. Carninci, P. et al. Genome-wide analysis of mammalian promoter architecture and evolution. Nat Genet 38, 626–35 (2006).

12. Nepal, C. et al. Dynamic regulation of the transcription initiation landscape at single nucleotide resolution during vertebrate embryogenesis. Genome Res 23, 1938–50 (2013).

13. Kawaji, H. et al. Comparison of CAGE and RNA-seq transcriptome profiling using clonally amplified and single-molecule next-generation sequencing. Genome Res 24, 708–17 (2014).

14. Haberle, V. et al. Two independent transcription initiation codes overlap on vertebrate core promoters. Nature 507, 381–5 (2014).

15. Nepal, C. et al. Dual-initiation promoters with intertwined canonical and TCT/TOP transcription start sites diversify transcript processing. Nat Commun 11, 168 (2020).

16. Chia, M. et al. High-resolution analysis of cell-state transitions in yeast suggests widespread transcriptional tuning by alternative starts. Genome Biol 22, 34 (2021).

17. Consortium, F. et al. A promoter-level mammalian expression atlas. Nature 507, 462–70 (2014).

18. Baranasic, D. et al. Multiomic atlas with functional stratification and developmental dynamics of zebrafish cis-regulatory elements. Nat Genet (2022).

19. Demircioglu, D. et al. A Pan-cancer Transcriptome Analysis Reveals Pervasive Regulation through Alternative Promoters. Cell 178, 1465–1477 e17 (2019).

20. Maunakea, A.K. et al. Conserved role of intragenic DNA methylation in regulating alternative promoters. Nature 466, 253–7 (2010).

21. Neri, F. et al. Intragenic DNA methylation prevents spurious transcription initiation. Nature 543, 72–77 (2017).

22. Meuleman, W. et al. Index and biological spectrum of human DNase I hypersensitive sites. Nature 584, 244–251 (2020).

23. Nepal, C. et al. Transcriptional, post-transcriptional and chromatin-associated regulation of pri-miRNAs, pre-miRNAs and moRNAs. Nucleic Acids Res 44, 3070–81 (2016).

24. Andersson, R. et al. An atlas of active enhancers across human cell types and tissues. Nature 507, 455–61 (2014).

25. Aran, D., Hu, Z. & Butte, A.J. xCell: digitally portraying the tissue cellular heterogeneity landscape. Genome Biol 18, 220 (2017).

26. Hoshida, Y. et al. Gene expression in fixed tissues and outcome in hepatocellular carcinoma. N Engl J Med 359, 1995–2004 (2008).

27. Subramanian, A. et al. Gene set enrichment analysis: a knowledge-based approach for interpreting genome-wide expression profiles. Proc Natl Acad Sci U S A 102, 15545–50 (2005).

28. Pinyol, R. et al. Molecular predictors of prevention of recurrence in HCC with sorafenib as adjuvant treatment and prognostic factors in the phase 3 STORM trial. Gut 68, 1065–1075 (2019).

29. Love, M.I., Huber, W. & Anders, S. Moderated estimation of fold change and dispersion for RNA-seq data with DESeq2. Genome Biol 15, 550 (2014).

30. Lin, D., Hiron, T.K. & O’Callaghan, C.A. Intragenic transcriptional interference regulates the human immune ligand MICA. EMBO J 37(2018).

31. Cinghu, S. et al. Intragenic Enhancers Attenuate Host Gene Expression. Mol Cell 68, 104–117 e6 (2017).

32. Saxonov, S., Berg, P. & Brutlag, D.L. A genome-wide analysis of CpG dinucleotides in the human genome distinguishes two distinct classes of promoters. Proc Natl Acad Sci U S A 103, 1412–7 (2006).

33. Illingworth, R.S. et al. Orphan CpG islands identify numerous conserved promoters in the mammalian genome. PLoS Genet 6, e1001134 (2010).

34. Roider, H.G., Lenhard, B., Kanhere, A., Haas, S.A. & Vingron, M. CpG-depleted promoters harbor tissue-specific transcription factor binding signals--implications for motif overrepresentation analyses. Nucleic Acids Res 37, 6305–15 (2009).

35. Cai, S.Y., Yu, D., Soroka, C.J., Wang, J. & Boyer, J.L. Hepatic NFAT signaling regulates the expression of inflammatory cytokines in cholestasis. J Hepatol 74, 550–559 (2021).

36. Bae, S. & Lesch, B.J. H3K4me1 Distribution Predicts Transcription State and Poising at Promoters. Front Cell Dev Biol 8, 289 (2020).

37. Soares, L.M. et al. Determinants of Histone H3K4 Methylation Patterns. Mol Cell 68, 773–785 e6 (2017).

38. Heintzman, N.D. et al. Distinct and predictive chromatin signatures of transcriptional promoters and enhancers in the human genome. Nat Genet 39, 311 – 8 (2007).

39. Hughes, A.L., Kelley, J.R. & Klose, R.J. Understanding the interplay between CpG island-associated gene promoters and H3K4 methylation. Biochim Biophys Acta Gene Regul Mech 1863, 194567 (2020).

40. Taniguchi, I., Iwaya, C., Ohnaka, K., Shibata, H. & Yamamoto, K. Genomewide DNA methylation analysis reveals hypomethylation in the low-CpG promoter regions in lymphoblastoid cell lines. Hum Genomics 11, 8 (2017).

41. Zilberman, D., Coleman-Derr, D., Ballinger, T. & Henikoff, S. Histone H2A.Z and DNA methylation are mutually antagonistic chromatin marks. Nature 456, 125–9 (2008).

42. Masalmeh, R.H.A. et al. De novo DNA methyltransferase activity in colorectal cancer is directed towards H3K36me3 marked CpG islands. Nat Commun 12, 694 (2021).

43. Jeziorska, D.M. et al. DNA methylation of intragenic CpG islands depends on their transcriptional activity during differentiation and disease. Proc Natl Acad Sci U S A 114, E7526–E7535 (2017).

44. Barski, A. et al. High-resolution profiling of histone methylations in the human genome. Cell 129, 823–37 (2007).

45. Edmunds, J.W., Mahadevan, L.C. & Clayton, A.L. Dynamic histone H3 methylation during gene induction: HYPB/Setd2 mediates all H3K36 trimethylation. EMBO J 27, 406–20 (2008).

46. Smolle, M. et al. Chromatin remodelers Isw1 and Chd1 maintain chromatin structure during transcription by preventing histone exchange. Nat Struct Mol Biol 19, 884–92 (2012).

47. Sen, P. et al. H3K36 methylation promotes longevity by enhancing transcriptional fidelity. Genes Dev 29, 1362–76 (2015).

48. Zhou, W. et al. DNA methylation loss in late-replicating domains is linked to mitotic cell division. Nat Genet 50, 591–602 (2018).

49. Partridge, E.C. et al. Occupancy maps of 208 chromatin-associated proteins in one human cell type. Nature 583, 720–728 (2020).

50. Karlsson, K., Lonnerberg, P. & Linnarsson, S. Alternative TSSs are coregulated in single cells in the mouse brain. Mol Syst Biol 13, 930 (2017).

51. Thomson, J.P. et al. CpG islands influence chromatin structure via the CpG-binding protein Cfp1. Nature 464, 1082–6 (2010).

52. Struhl, K. Transcriptional noise and the fidelity of initiation by RNA polymerase II. Nat Struct Mol Biol 14, 103–5 (2007).

53. Ayoubi, T.A. & Van De Ven, W.J. Regulation of gene expression by alternative promoters. FASEB J 10, 453–60 (1996).

54. Langmead, B., Trapnell, C., Pop, M. & Salzberg, S.L. Ultrafast and memory-efficient alignment of short DNA sequences to the human genome. Genome Biol 10, R25 (2009).

55. Li, H. et al. The Sequence Alignment/Map format and SAMtools. Bioinformatics 25, 2078–9 (2009).

56. Amemiya, H.M., Kundaje, A. & Boyle, A.P. The ENCODE Blacklist: Identification of Problematic Regions of the Genome. Sci Rep 9, 9354 (2019).

57. Gao, T. & Qian, J. EnhancerAtlas 2.0: an updated resource with enhancer annotation in 586 tissue/cell types across nine species. Nucleic Acids Res 48, D58–D64 (2020).

58. Lee, B.T. et al. The UCSC Genome Browser database: 2022 update. Nucleic Acids Res 50, D1115–D1122 (2022).

59. Huang, H. et al. Histone H3 trimethylation at lysine 36 guides m(6)A RNA modification co-transcriptionally. Nature 567, 414–419 (2019).

60. Ramirez, F. et al. deepTools2: a next generation web server for deep-sequencing data analysis. Nucleic Acids Res 44, W160–5 (2016).

61. Goldman, M.J. et al. Visualizing and interpreting cancer genomics data via the Xena platform. Nat Biotechnol 38, 675–678 (2020).

62. Consortium, E.P. An integrated encyclopedia of DNA elements in the human genome. Nature 489, 57–74 (2012).

63. Quinlan, A.R. & Hall, I.M. BEDTools: a flexible suite of utilities for comparing genomic features. Bioinformatics 26, 841–2 (2010).

64. Heinz, S. et al. Simple combinations of lineage-determining transcription factors prime cis-regulatory elements required for macrophage and B cell identities. Mol Cell 38, 576–89 (2010).

65. Liu, J. et al. An Integrated TCGA Pan-Cancer Clinical Data Resource to Drive High-Quality Survival Outcome Analytics. Cell 173, 400–416 e11 (2018).

